# Age Differences in Hippocampal Neural Timescales During Movie-Viewing

**DOI:** 10.64898/2026.05.05.723065

**Authors:** Nichole R. Bouffard, Angelique I. Delarazan, Ata B. Karagoz, Jeffrey M. Zacks, Zachariah M. Reagh

## Abstract

Episodic memory requires integrating information across multiple scales, a process theorized to be supported by a gradient of neural timescales along the anterior-posterior axis of the hippocampus that enables both coarse-and fine-grained representations. Aging is associated with changes in hippocampal function and declines in fine-grained episodic memory, but whether this impacts the gradient organization of the hippocampus is unknown. Additionally, the relationship between the neural timescales of the hippocampus and memory specificity remains unclear. Here, we analyzed the length of timescales of individual voxels in the hippocampus during movie-viewing, along with subsequent recall data, in a sample of young and older participants. Younger adults showed the expected anterior-to-posterior timescale gradient, replicating prior work. In contrast, older adults exhibited a reversal of the expected gradient. Older adults’ recall was coarser and more gist-like than that of younger adults. In younger adults, longer neural timescales were associated with less specific, more gist-like recall; this was seen predominantly in the posterior-lateral hippocampus. In contrast, no relationship between neural timescales and recall were observed in older adults. An exploratory analysis revealed a similar relationship between neural timescales and memory specificity in cortical regions, in younger but not older adults. These findings suggest that aging alters the organization of neural activity throughout the hippocampus and that neural timescales in the hippocampus and cortex are related to the specificity of memory.

**Significance statement:** As people age, episodic memories become more gist-like and less detailed. The hippocampus, which supports both gist and detailed memory, exhibits a neural timescale gradient—from slow-changing activity (longer timescales) to fast-changing activity (shorter timescales). This organization is theorized to support coarse-and fine-grained memory, respectively, yet a direct link to the age-related shift towards gist-like memory remains unestablished. Here, we identify an age-related shift in the hippocampal timescale gradient that parallels a decline in memory specificity. Furthermore, longer timescales in the hippocampus and cortical regions correlated with decreased memory specificity in younger adults. These findings demonstrate that aging is associated with a reorganization of hippocampal activity and that cortical timescales during encoding may relate to the specificity of memory.

## Introduction

Aging is associated with a shift in episodic memories toward “gist” and a reduction in retrieved details (Delarazan et al., 2023; Greene & Naveh-Benjamin, 2020; Grilli & Sheldon, 2022; Levine et al., 2002). However, despite decades of investigation, the neural mechanisms underlying such a shift remain unclear. Current theoretical frameworks posit that the hippocampus supports a hierarchy of memory specificity through a functional gradient along its long axis, transitioning from coarse, global representations in the anterior hippocampus to fine, local representations in the posterior hippocampus (Evensmoen et al., 2015; Fanselow & Dong, 2010; Grady, 2020; Milivojevic & Doeller, 2013; Poppenk et al., 2013a; Strange et al., 2014). Recently, this representational gradient has been linked to a gradient of intrinsic neural dynamics in fMRI, where the activity of single voxels in the anterior hippocampus is stable over long temporal windows and the activity of single voxels in the posterior hippocampus changes more rapidly (Bouffard, Golestani, et al., 2023; Brunec, Bellana, et al., 2018; Coughlan et al., 2023; Raut et al., 2020). Importantly, this gradient has been related to spatial navigation (Bouffard, Golestani, et al., 2023) and disruptions to the gradient in temporal lobe epilepsy have been related to memory deficits (Bouffard et al., 2026). This physiological architecture ideally situates the hippocampus to support the simultaneous processing of gist and detailed memories.

It is possible that changes to functional gradients in the hippocampus underlie age-related shifts toward gist memory. Aging is known to affect hippocampal structure (De Leon et al., 1997; Golomb et al., 1993), task-related activation (Cabeza, Grady, et al., 1997; Cabeza, McIntosh, et al., 1997), and functional connectivity to memory networks (Andrews-Hanna et al., 2007; Damoiseaux et al., 2016; Grady et al., 2003; Stark et al., 2021). Furthermore, age-related cognitive decline has been linked to widespread changes in neural signal dynamics (Garrett et al., 2011, 2013) and to slowing of neural state transitions (Lugtmeijer et al., 2025). Such a disruption to representational timescales of activity in the hippocampus could have downstream consequences relating to temporal dynamics across the whole brain (Irish & Vatansever, 2020; Raut et al., 2020; Stephens et al., 2013), possibly disrupting hierarchies of representational timescales from sensory regions to higher-order association cortex (Hasson et al., 2008; Honey et al., 2012).

Here, we tested whether the distinct anterior-posterior gradient found in younger adults is diminished or altered in older adults. We further predicted that changes to the gradient relate to a reduction in memory specificity in older adults relative to younger adults. To test this, we used a movie-viewing paradigm, which goes beyond prior work that has relied primarily on resting-state fMRI. Movie-viewing offers a significant advantage over traditional lab-based tasks because it requires the integration of information across multiple temporal scales, more closely approximating the demands of everyday episodic encoding where older adults often demonstrate a shift toward coarse, gist-like memory. This approach allowed us to determine whether the hippocampal gradients previously identified during resting state are also maintained during continuous, naturalistic processing. While we primarily focused on the hippocampus, we also conducted exploratory analyses to determine if the relationship between neural timescales and memory specificity is a unique computational feature of this region or if it extends to broader cortical control areas.

We report two key findings that challenge and expand our understanding of these dynamics. First, the canonical hippocampal gradient observed in younger adults is not merely attenuated but *reversed* in older adults, suggesting a profound reorganization of temporal processing in the older brain. Second, we observe a positive relationship between hippocampal neural timescales and memory specificity in younger adults. However, contrary to a hippocampus-exclusive hypothesis, we found that this relationship extends to control regions, including the visual and motor cortices. These results suggest that the link between neural timescales and memory specificity is likely a signature of a broader brain state during encoding, rather than a process confined solely to the hippocampus.

## Methods

### Participants

Younger adult participants (N = 39, age range = 18-33) were recruited from the University of California, Davis Psychology experiment website (Sona Systems). Older adult participants (N = 26, age range = 60-84) were recruited from the community via flyer and email advertisement. All participants were paid $20 per hour and gave written informed consent in accordance with the University of California, Davis Institutional Review Board. Exclusion criteria included magnetic bodily implants, claustrophobia, a history of major psychiatric or neurological disorders, a concussion in the past 6 months, untreated diabetes or hypertension, current drug or alcohol abuse, uncorrected hearing, uncorrected vision, or left-handedness.

### Stimuli

The video participants watched was an episode of Curb Your Enthusiasm (Season 1, Episode 7). The beginning and ending credits were trimmed from the episode, resulting in a 26-minute video. The video was presented to participants in the MRI scanner using PsychoPy version 3.1.5, and the stimulus onset was synchronized with MRI pulse sequences via fiber optic trigger pulses sent to the stimulus computer. After watching the video, participants completed a resting-state scan and performed verbal recall of the entire video while in the scanner (see Procedure).

### Procedure

Scanning took place in a Siemens 3 T Skyra with a 32-channel head coil. Participants were fitted with MRI-compatible earbuds with replaceable foam inserts (MRIaudio) and were provided with additional foam padding inside the head coil to protect hearing and to mitigate head motion. Participants were additionally given body padding, blankets, and corrective lenses as needed. An MRI compatible microphone (Optoacoustics FOMRI-III) with bidirectional noise-canceling was affixed to the head coil, and the receiver (covered by a disposable sanitary pop screen) was positioned over the participant’s mouth.

High-resolution T1-weighted structural images were acquired using a magnetization prepared rapid acquisition gradient echo (MPRAGE) pulse sequence (FOV = 256mm, image matrix = 256 x 256, sagittal slices = 208, thickness = 1mm). Functional images were acquired using a multi-band gradient echo planar imaging (EPI) sequence (TR = 1220ms, TE = 24ms, FOV = 192mm, image matrix = 64 x 64, flip angle = 67, multi-band factor = 2, axial slices = 38, voxel size = 3mm isotropic, P » A phase encoding, AC-PC alignment). A single four-TR functional scan of reverse phase encoding polarity (A » P) was acquired for unwarping. The video watching functional scans were approximately 26 minutes long, consisting of 1278 volumes.

After the movie-viewing task, an 8-minute resting state fMRI scan was collected. During this time participants were instructed to keep as still as possible, relax, and think about anything they’d like. After the resting state scan, participants completed a third and final functional scan where a fixation cross was presented on the screen and participants were told to recall the entire episode, from beginning to end, in as much detail as possible. Their recall was recorded using an MRI-compatible microphone. We were primarily interested in hippocampal neural timescales during encoding and how these encoding neural signatures related to subsequent recall performance, therefore analyses of the resting-state and recall scans are not included in the present manuscript but will be included in a forthcoming paper (Delarazan, Ranganath, et al., 2026). After the scan, older adults completed a series of neuropsychological tests.

### fMRI Data Preprocessing

Results included in this manuscript come from preprocessing performed using *fMRIPrep* 20.2.7 (Esteban, Blair, et al., 2018; Esteban, Markiewicz, et al., 2018); RRID:SCR_016216), which is based on *Nipype* 1.7.0 (K. Gorgolewski et al., 2011; K. J. Gorgolewski et al., 2018).

#### Anatomical Data Preprocessing

A total of 1 T1-weighted (T1w) images were found within the input BIDS dataset. The T1-weighted (T1w) image was corrected for intensity non-uniformity (INU) with N4BiasFieldCorrection (Tustison et al., 2010), distributed with ANTs 2.3.3 (Avants et al., 2008), and used as T1w-reference throughout the workflow. The T1w-reference was then skull-stripped with a *Nipype* implementation of the antsBrainExtraction.sh workflow (from ANTs), using OASIS30ANTs as target template. Brain tissue segmentation of cerebrospinal fluid (CSF), white-matter (WM) and gray-matter (GM) was performed on the brain-extracted T1w using fast (FSL 5.0.9, RRID:SCR_002823, (Zhang et al., 2001). Brain surfaces were reconstructed using recon-all (FreeSurfer 6.0.1, RRID:SCR_001847, (Dale et al., 1999)), and the brain mask estimated previously was refined with a custom variation of the method to reconcile ANTs-derived and FreeSurfer-derived segmentations of the cortical gray-matter of Mindboggle (RRID:SCR_002438, (Klein et al., 2017). Volume-based spatial normalization to two standard spaces (MNI152NLin2009cAsym, MNI152Lin) was performed through nonlinear registration with antsRegistration (ANTs 2.3.3), using brain-extracted versions of both T1w reference and the T1w template. The following templates were selected for spatial normalization: *ICBM 152 Nonlinear Asymmetrical template version 2009c* [(Fonov et al., 2009); RRID:SCR_008796; TemplateFlow ID: MNI152NLin2009cAsym], *Linear ICBM Average Brain (ICBM152) Stereotaxic Registration Model* [(Mazziotta et al., 1995) TemplateFlow ID: MNI152Lin],

#### Functional Data Preprocessing

For each of the 3 BOLD runs found per subject (across all tasks and sessions), the following preprocessing was performed. First, a reference volume and its skull-stripped version were generated using a custom methodology of *fMRIPrep*. A B0-nonuniformity map (or *fieldmap*) was estimated based on two (or more) echo-planar imaging (EPI) references with opposing phase-encoding directions, with 3dQwarp Cox and Hyde (1997) (AFNI 20160207). Based on the estimated susceptibility distortion, a corrected EPI (echo-planar imaging) reference was calculated for a more accurate co-registration with the anatomical reference. The BOLD reference was then co-registered to the T1w reference using bbregister (FreeSurfer) which implements boundary-based registration (Greve & Fischl, 2009). Co-registration was configured with six degrees of freedom. Head-motion parameters with respect to the BOLD reference (transformation matrices, and six corresponding rotation and translation parameters) are estimated before any spatiotemporal filtering using mcflirt (FSL 5.0.9, (Jenkinson et al., 2002). BOLD runs were slice-time corrected to 0.568s (0.5 of slice acquisition range 0s-1.14s) using 3dTshift from AFNI 20160207 (Cox & Hyde, 1997)RRID:SCR_005927). The BOLD time-series were resampled onto the following surfaces (FreeSurfer reconstruction nomenclature): *fsaverage*. The BOLD time-series (including slice-timing correction when applied) were resampled onto their original, native space by applying a single, composite transform to correct for head-motion and susceptibility distortions. These resampled BOLD time-series will be referred to as *preprocessed BOLD in original space*, or just *preprocessed BOLD*. The BOLD time-series were resampled into several standard spaces, correspondingly generating the following *spatially-normalized, preprocessed BOLD runs*: MNI152NLin2009cAsym, MNI152Lin. First, a reference volume and its skull-stripped version were generated using a custom methodology of *fMRIPrep*. Several confounding time-series were calculated based on the *preprocessed BOLD*: framewise displacement (FD), DVARS and three region-wise global signals. FD was computed using two formulations following Power (absolute sum of relative motions, (Power et al., 2014)) and Jenkinson (relative root mean square displacement between affines, (Jenkinson et al., 2002))). FD and DVARS are calculated for each functional run, both using their implementations in *Nipype* (following the definitions by (Power et al., 2014). The three global signals are extracted within the CSF, the WM, and the whole-brain masks. Additionally, a set of physiological regressors were extracted to allow for component-based noise correction (*CompCor*, (Behzadi et al., 2007). Principal components are estimated after high-pass filtering the *preprocessed BOLD* time-series (using a discrete cosine filter with 128s cut-off) for the two *CompCor* variants: temporal (tCompCor) and anatomical (aCompCor). tCompCor components are then calculated from the top 2% variable voxels within the brain mask. For aCompCor, three probabilistic masks (CSF, WM and combined CSF+WM) are generated in anatomical space. The implementation differs from that of Behzadi et al. in that instead of eroding the masks by 2 pixels on BOLD space, the aCompCor masks are subtracted a mask of pixels that likely contain a volume fraction of GM. This mask is obtained by dilating a GM mask extracted from the FreeSurfer’s *aseg* segmentation, and it ensures components are not extracted from voxels containing a minimal fraction of GM. Finally, these masks are resampled into BOLD space and binarized by thresholding at 0.99 (as in the original implementation). Components are also calculated separately within the WM and CSF masks. For each CompCor decomposition, the *k* components with the largest singular values are retained, such that the retained components’ time series are sufficient to explain 50 percent of variance across the nuisance mask (CSF, WM, combined, or temporal). The remaining components are dropped from consideration. The head-motion estimates calculated in the correction step were also placed within the corresponding confounds file. The confound time series derived from head motion estimates and global signals were expanded with the inclusion of temporal derivatives and quadratic terms for each (Satterthwaite et al., 2013). Frames that exceeded a threshold of 0.5 mm FD or 1.5 standardised DVARS were annotated as motion outliers. All resamplings can be performed with *a single interpolation step* by composing all the pertinent transformations (i.e. head-motion transform matrices, susceptibility distortion correction when available, and co-registrations to anatomical and output spaces). Gridded (volumetric) resamplings were performed using antsApplyTransforms (ANTs), configured with Lanczos interpolation to minimize the smoothing effects of other kernels (Lanczos, 1964). Non-gridded (surface) resamplings were performed using mri_vol2surf (FreeSurfer).

Many internal operations of *fMRIPrep* use *Nilearn* 0.6.2 (Abraham et al., 2014, RRID:SCR_001362), mostly within the functional processing workflow. For more details of the pipeline, see <u>the section corresponding to workflows in *fMRIPrep*’s</u> <u>documentation</u>.

#### Copyright Waiver

The above boilerplate text was automatically generated by fMRIPrep with the express intention that users should copy and paste this text into their manuscripts *unchanged*. It is released under the <u>CC0</u> license.

After running fMRIPrep, we conducted additional preprocessing steps. Global signal, CSF and white matter regressors from the fMRIprep output were regressed out using *Nilearn 0.10.4* (Abraham et al., 2014) and *Nibabel 5.3.1* (Brett et al., 2024) in custom python scripts. Linear detrending and bandpass temporal filtering (0.1-0.01 Hz) were applied. Our main measure of interest, intrinsic neural timescale, is dependent on the autocorrelative structure of the timeseries data, therefore motion and motion spikes are of particular concern. Because older adults tend to have more motion during scans, we took a conservative approach to control for motion confounds. We excluded participants who had a mean framewise displacement greater than 0.7 mm. All 24 motion confounds plus their derivatives (computed by fMRIprep) were regressed out.

Motion spikes are particularly disruptive to the continuity of the timeseries and the effect that this censoring has on the intrinsic neural timescale measure is still unknown. While prior studies used different approaches to mitigate motion spikes (Bouffard et al., 2023, 2026; Raut et al., 2020; Xie et al., 2023), there has been no comparison of how motion scrubbing affects the intrinsic neural timescale values. We therefore conducted all our analyses on both the non-scrubbed data (but with the same participant exclusion criteria and previous preprocessing steps applied) and the motion scrubbed data. The results presented in the main text are from the scrubbed dataset. We present our analyses of the non-scrubbed data in the Supplemental materials (Figures S1 and S2).

To identify motion spikes in the data we used the approach outlined by Wilford and colleagues (2025). We first temporally filtered the FWD and stdDVars fMRIprep confounds using the same temporal filter we applied to the data (0.1-0.01Hz). We then identified motion spikes as any TRs that were FWD > 0.5 mm or stdDvars > 1.5. There were three younger adults and three older adults who had excessive motion spikes (>250) and therefore were excluded from our analyses (both scrubbed and non-scrubbed analyses). To preserve the autocorrelative structure of the data and to prevent an artificial inflation of the autocorrelation by interpolating, we replaced flagged TRs with NAs as placeholders and then computed the lagged autocorrelation structure of the data over successive single TR lag shifts. This motion spike censoring step was conducted concurrently with the calculation of the autocorrelation using a custom Python script (https://github.com/atakaragoz/fmri_autocorr).

#### Participant Exclusions

We excluded one younger adult and one older adult participant who had mean framewise displacements greater than 0.7 mm. We also excluded three younger and three older adults who had excessive motion spikes (operationalized as >250 TRs). Additional exclusions included two younger adults who had significant signal drop out, one younger adult who fell asleep during the encoding scan, and two younger adults whose data couldn’t be accessed due corruption of the relevant files. The final sample consisted of of 30 younger adults (*M_age_* = 22.33, *SD_age_* = 2.77, Female N = 21) and 22 older adults (*M_age_* = 68.52, *SD_age_* = 6.81, Female N = 11). Due to technical issues with the microphone in the scanner during data collection, three older adults did not have recall audio files that could be transcribed; therefore, analyses of the behavioral recall data are from a final sample of 19 older adults.

#### fMRI Data Analysis – Intrinsic Neural Timescales (INTs)

After preprocessing, we used the right and left hippocampal mask from the Harvard-Oxford Atlas to mask only hippocampal voxels. We then computed the intrinsic neural timescale of each individual voxel (Raut et al., 2020; Watanabe et al., 2019; Xie et al., 2023). The intrinsic neural timescale, or INT, for a given voxel is defined as the area under the curve of the autocorrelation function (ACF) during the initial positive period, also called ACW0 (i.e., the autocorrelation window of positive values before the function reaches 0). INTs were computed using the following steps: First, the autocorrelation function (ACF) of the fMRI signal of each voxel was calculated by computing successive lagged-shifted correlations of the BOLD signal with itself (e.g., AC_lag1_, AC_lag2_, AC_lag3_, etc.) (Figure 1B). The ACF values for in the initial period where the ACF is positive were summed and multiplied by the repetition time (TR) (i.e., the area under the curve). Although we report the average AC_lag1_, ACW0, and INTs for each cluster and each group (Table S2), our primary analyses used INTs as our measure of interest. We Z-scored the INTs values within each participant for the plots to better visualize comparisons between anterior-medial and posterior-lateral INTs.

**Figure 1.**
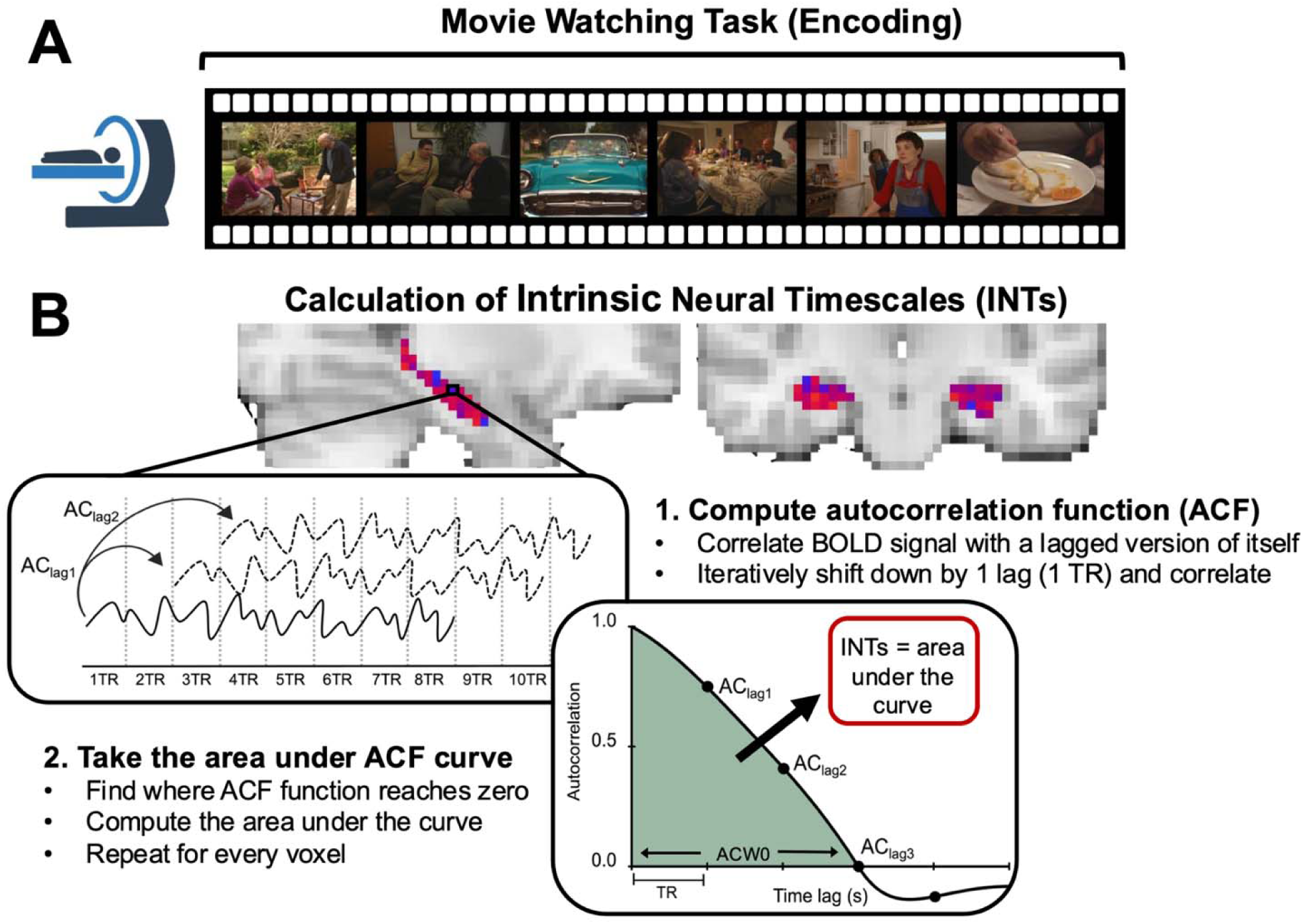
**A. fMRI Paradigm.** Younger and older adult participants were scanned while watching a 26-minute-long episode of Curb your Enthusiasm. After watching the episode, participants were instructed to recall aloud everything they could remember from the episode, in as much detail as possible. **B. Calculation of Intrinsic Neural Timescales.** For each voxel in the hippocampus, the autocorrelation function (ACF) was computed by calculating correlations of the signal with a lagged version of itself. For lagged correlations, the signal is successively shifted by a lag of 1 TR (or 1.22 seconds). The area under the curve of the autocorrelation function is computed for the window in which the ACF was positive, or ACW0 (i.e., window before it reaches zero). The area under the curve is the INTs, or intrinsic neural timescale.

#### Hippocampal ROIs

One of our primary aims was to investigate whether the neural timescale gradient observed in resting state fMRI (Bouffard, Golestani et al., 2023) is also found during movie-viewing. Additionally, we investigated whether this gradient is related to aging and whether both younger and older adults demonstrate a similar gradient. In our previous work, we measured hippocampal neural timescales in a sample of 44 younger adults from the Human Connectome Project database. Using a data-driven approach, we found that neural timescales during resting state fMRI were organized in a gradient, such that voxels with longer timescales clustered in the anterior-medial hippocampus and voxels with shorter timescales clustered in the posterior-lateral hippocampus (Bouffard, Golestani et al., 2023). We used these previously defined, data-driven clusters as a means of unbiased ROI selection applied to the present dataset.

#### Recall Scoring

Each participant’s recall was manually scored for gist and detail by comparing participant recall to the transcript of the episode. The episode was divided into meaningful event units (Sargent et al., 2013) which resulted in 37 possible gist event units and 195 detail event units for each event (Delarazan, March, et al., 2026). The audio recordings of each participants’ recall were transcribed to text using Whisper (*Openai-Whisper*, n.d.). Due to technical issues with the microphone in the scanner, three older adults did not have audio files that could be transcribed. The analyses therefore include data from 30 younger adults and 19 older adults. Two independent raters (E.K. and H.C.) segmented and labeled each participant’s recall, matching every recalled event to the specific scene in the episode it pertained to. Each event was annotated with a structured description specifying who was involved, where it occurred, and what happened. The raters scored each recalled event for specificity by awarding gist units when a participant conveyed the general meaning of an event description and detail units when the participant additionally recalled specific components (who/where/what). Detail scoring was nested within gist such that any detail unit necessarily implied the corresponding gist unit, whereas gist units could be awarded in the absence of detail. The intraclass correlation coefficient (ICC) was calculated to assess the agreement between two raters. The analysis revealed an excellent level of agreement. For single raters using a fixed-effects model (ICC3), the ICC was 0.96, 95% CI[0.77,1.0], F(5,5)=56.09, p<.001. For average raters using the same model (ICC3k), the ICC was 0.98, 95% CI[0.87,1.0], F(5,5)=56.09, p<.001.

The specificity ratio was calculated to quantify the proportion of gist and detail in each participant’s recall. For each participant, the number of detail ratings was subtracted from the number of gist ratings. This was then divided by the total number of ratings (gist + detail). The formula is below:

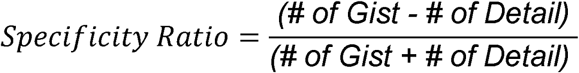

Specificity ratio scores range between 0 and 1, where a value of 0 represents maximal number of details recalled, and a value of 1 represents only gist-level recall and no details.

#### Neuropsychological Testing

At the end of the experiment, older participants completed a battery of neuropsychological tests to evaluate cognitive impairments that might be relevant for assessing task performance. These tests included: Craft21 Recall Immediate, Craft21 Recall Delayed, Montreal Cognitive Assessment (MoCA), and Multilingual Naming Test (MINT). Briefly, Craft21 assesses recall for narratives (Weintraub et al., 2018), MoCA coarsely assesses cognitive ability (Nasreddine et al., 2005), and MINT assesses the ability to name objects in English (Gollan et al., 2012). The summary of scores are reported in Supplemental Table S1.

## Results

### Gradients of Hippocampal INTs Differ with Age

We analyzed the movie-viewing fMRI data from younger (N=30) and older adults (N=22). Previous work found high INTs in the anterior-medial hippocampus and low INTs in the posterior-lateral hippocampus among younger adults (Bouffard et al., 2023; Coughlan et al., 2023). Here, we tested whether the gradient of INTs in the hippocampus previously observed in younger adults is similar in older adults. We used hippocampal ROIs from Bouffard, Golestani et al. (2023) to mask the hippocampus and label voxels as either anterior-medial or posterior-lateral clusters (Figure 2A). We report two analyses: one at the level of the cluster (averaging INTs within each cluster) and one at the level of the voxel (examining INTs within each voxel).

**Figure 2.**
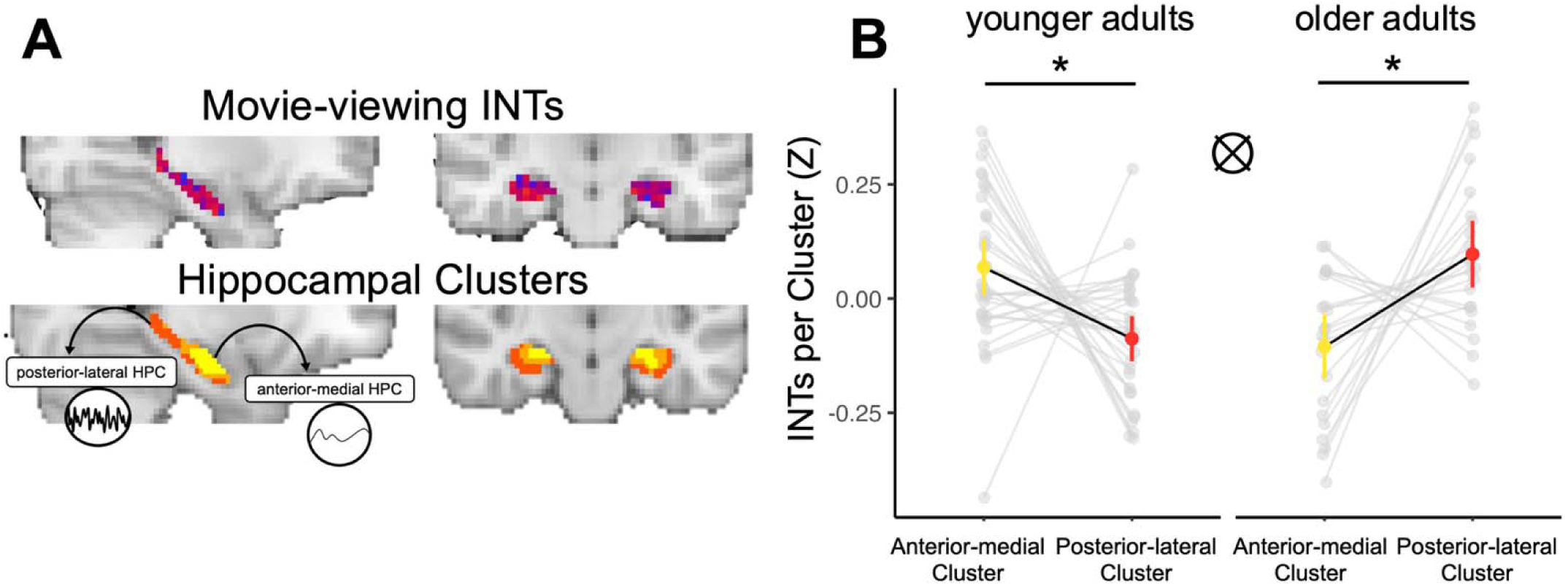
**A. Example map of hippocampal INTs during movie-viewing and hippocampal clusters.** INTs were computed for every voxel in the hippocampus during movie-viewing. Data-driven clusters from an independent dataset were used to mask the movie-viewing INTs data (Bouffard, Golestani, et al., 2023). In previous studies with younger adults, voxels within the anterior-medial cluster tend to have longer neural timescales and voxels in posterior-lateral cluster tend to have shorter neural timescales. Clusters were used to group and average the INTs data into anterior-medial and posterior-lateral clusters. **B. Hippocampal INTs gradient in younger and older adults.** The average INTs in the anterior-medial cluster is depicted in yellow and the average INTs in the posterior-lateral cluster is depicted in red. INTs values were z-scored within each participant to better visualize comparisons between anterior-medial and posterior-lateral INTs. Error bars represent 95% CIs. Individual participant scores are depicted in grey. There was a significant interaction between age group and cluster, such that younger adults demonstrated the expected gradient, where INTs in the anterior-medial cluster were significantly greater than INTs in the posterior-lateral cluster. Older adults, however, demonstrated a reversal of this gradient, such that the anterior-medial cluster had lower INTs than the posterior-lateral cluster, * p < 0.05.

We first ran a linear mixed effects model on the average INTs within each cluster and included fixed effect predictors for hemisphere (Left, Right), age group (younger, older), cluster (anterior-medial, posterior-lateral). The random effects structure was determined by first computing the maximal model justified by the experimental design (Barr et al., 2013), including participant and age group as random intercepts and all the associated random slopes. We then systematically trimmed the random intercepts and random slopes from the random effects structure until the model converged, while avoiding a singular solution (Singmann & Kellen, 2019). The resulting model included participant as the only random intercept and cluster as the only random slope in the random effects term. The full model formula: average INTs per cluster ∼ hemisphere * age group * cluster + (1 + cluster|participant). For post hoc tests of the main effects and interactions, the degrees of freedom was calculated using Kenward-Rogers method and p values were adjusted for multiple comparisons using the Tukey method.

We did not find significant effects of hemisphere, age group, or cluster. However, we found a significant interaction between age group and cluster (F(1,50) = 15.45, p < 0.001) (Figure 2B). Parsing the interaction between age group and cluster revealed that the INTs in the anterior-medial cluster were greater than the posterior-lateral cluster in younger adults (t(50) = 2.82, p = 0.033), in line with prior work. In older adults this was reversed, and the INTs in the posterior-lateral cluster were greater than the anterior-medial cluster (t(50) = 2.75, p = 0.039). These results suggest that, although there is no significant difference in overall hippocampal INTs between older (M = 2.30, SD =0.19) and younger adults (M = 2.36, SD = 0.25), younger adults demonstrate the expected anterior-medial to posterior-lateral gradient of INTs whereas older adults demonstrate a complete reversal of this gradient. Analysis of the non-scrubbed data yielded a similar pattern, but older adults did not show a significant difference in INTs between the anterior-medial cluster and posterior-lateral cluster, although numerically the anterior-medial cluster was lower than the posterior-lateral cluster (see Figures S1).

We repeated this analysis at the level of the individual voxel, using the INTs per voxel as the dependent variable (instead of the average INTs per cluster). We ran a linear mixed effects model on the INT for each voxel and included fixed effect predictors for hemisphere (Left, Right), age group (Younger, Older), and resting state cluster (anterior-medial, posterior-lateral). The full model formula: INTs per voxel ∼ hemisphere * age group * cluster + (1 + cluster|participant). For post hoc tests of the main effects and interactions, degrees of freedom were calculated using an asymptotic method (reporting z ratio instead of t statistic) and p values were adjusted for multiple comparisons using the Tukey method.

We did not find a significant main effect of hemisphere, age group, or cluster. We found a significant interaction between age group and cluster (F(1,50.01) = 15.55, p < 0.001). Analysis of the interaction between age group and cluster revealed that the INTs in the anterior-medial cluster were greater than the posterior-lateral cluster in younger adults (z = 2.84, p = 0.023). In older adults, this was reversed such that INTs in the posterior-lateral cluster were greater than the anterior-medial (z = 2.76, p = 0.029). There was also a significant interaction between hemisphere and age group (F(1, 12527.01) = 4.97, p = 0.026). However, none of the pairwise comparisons were significant (ps > 0.17). When analyzing INTs at the individual voxel level, we again find that younger adults demonstrate the expected anterior-medial to posterior-lateral gradient while older adults demonstrate a complete reversal of this gradient.

### Relationship Between Hippocampal INTs and Recall Specificity

#### Older adults have more gist-like recall

We scored participant’s recall of the TV episode for gist and detail and computed the specificity ratio. Higher values indicated more gist-like recall and lower values indicate more detailed recall. We found that older adults (N = 19) had higher specificity ratio scores compared to younger adults (N = 30) (M_older_ = 0.40, SD_older_ = 0.15; M_younger_ = 0.24, SD_younger_ = 0.18; t(43.75) = 3.27, p = 0.002; Figure 3A). In line with previous studies, this suggests that older adults had more gist-like recall of the episode and less detailed recall compared to younger adults (Delarazan et al., 2023; Greene & Naveh-Benjamin, 2020; Grilli & Sheldon, 2022).

**Figure 3.**
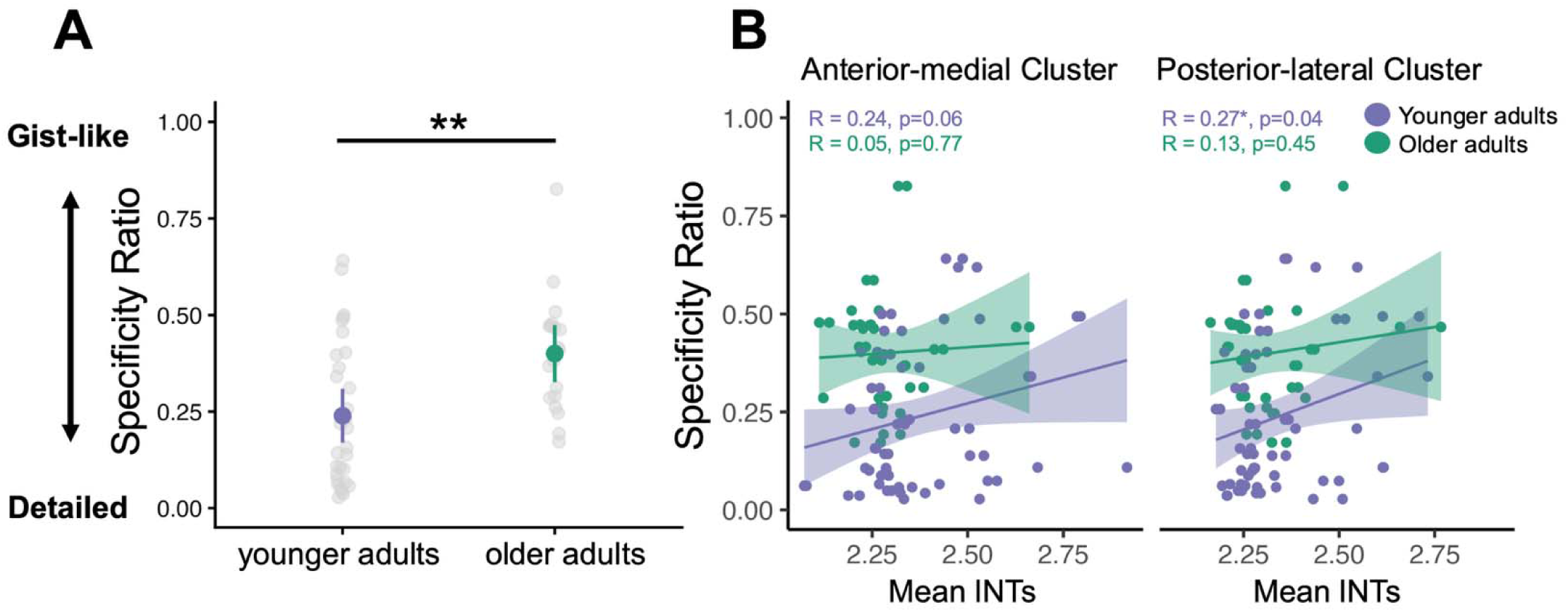
**A. Recall specificity for younger and older adults.** Participant recall was scored by tabulating the number of gist and detail units in each participant’s recall and then the specificity ratio was calculated by taking the difference between the number of gist and detail units divided by the sum of the total number of gist and detail units. The group mean specificity score for younger adults is depicted in purple and the group mean for older adults is depicted in green. Error bars represent the 95% CIs. Individual participant scores are depicted in grey. Older adults had significantly greater specificity ratio scores compared to younger adults, suggesting that older adults had more gist-like recall and less detailed recall than younger adults, ** p < 0.01. **B. Correlation between specificity ratio and mean INTs in the anterior-medial and posterior-lateral clusters.** Pearson’s correlations were computed between the specificity ratio and mean INTs within each cluster. Each point represents the mean INTs an individual participant and hemisphere. Solid lines indicate the linear regression fit; shaded ribbons indicate 95% CIs. There was a positive relationship, where longer timescales were related to more gist-like recall in the posterior-lateral cluster in younger but not older adults. While we did find a significant correlation in the posterior-lateral cluster, none of the correlations survived FDR multiple comparisons correction.

#### Longer timescales in the hippocampus during movie-viewing are related to gist-like recall in younger adults

To investigate how neural timescales in the hippocampus during movie-viewing are related to memory specificity, we correlated hippocampal INTs and specificity ratio scores. First, the INTs for was computed for each voxel. The INTs was then averaged across voxels in each cluster in the left and right hippocampus separately. Average INTs per cluster was correlated with the specificity scores using Pearson’s correlation (Figure 3B). We applied FDR correction for multiple comparisons and report both the uncorrected and corrected p values.

We found a positive correlation between mean INTs and specificity scores in the posterior-lateral cluster in younger adults (R = 0.27, p = 0.04, p_corrected_ = 0.13). However, we note that this result does not remain significant upon correcting for multiple comparisons (i.e., other correlations examined). We did not find a significant correlation in the anterior-medial cluster (R = 0.24, p = 0.06, p_corrected_ = 0.13). We did not find any relationship between INTs and specificity ratio scores in either cluster in older adults (Anterior-medial: R = 0.05, p = 0.77, p_corrected_ = 0.77; Posterior-lateral: R = 0.13, p = 0.45, p_corrected_ = 0.61). The positive relationship between mean INTs and specificity scores in younger adults suggests that greater hippocampal INTs, or relatively more “sluggish” signal, in the posterior-lateral cluster during encoding is associated with more gist-like recall. It is important to note that these correlations did not survive multiple comparisons correction, perhaps due to low power. Nonetheless, in contrast to younger adults, no evidence for any relationship between INTs and memory specificity was observed in older adults. Similar results were observed in analyses of the non-scrubbed data (see Supplemental materials Figure S2).

### INTs are Related to Recall Specificity in Cortical ROIs

We conducted exploratory analyses to examine two potential possibilities for the relationship between neural timescales and episodic memory: (1) The neural timescale gradient is a unique computational feature of the hippocampus and that its relationship to memory specificity is exclusive to this region. (2) The relationship between neural timescales and memory specificity reflects a broader, brain-wide mechanism. In the second view, differences in neural timescales observed in the hippocampus may follow a global brain state that facilitates either detailed or gist-like retention rather than a localized computation.

To investigate whether the relationship between INTs and memory specificity was specific and unique to the hippocampus or whether this relationship is also found in other areas of the brain, we computed the correlation between INTs and specificity scores in two controls regions: the early visual cortex and motor cortex. We chose these regions because they are associated with perception and sensory representations and we did not have strong predictions that these regions would relate memory specificity. We computed the averaged the INTs across voxels in the bilateral early visual cortex and motor cortex. Mean INTs per ROI was correlated with the specificity scores using Pearson’s correlation (Figure 4). We applied FDR correction for multiple comparisons and report both the uncorrected and corrected p values.

**Figure 4.**
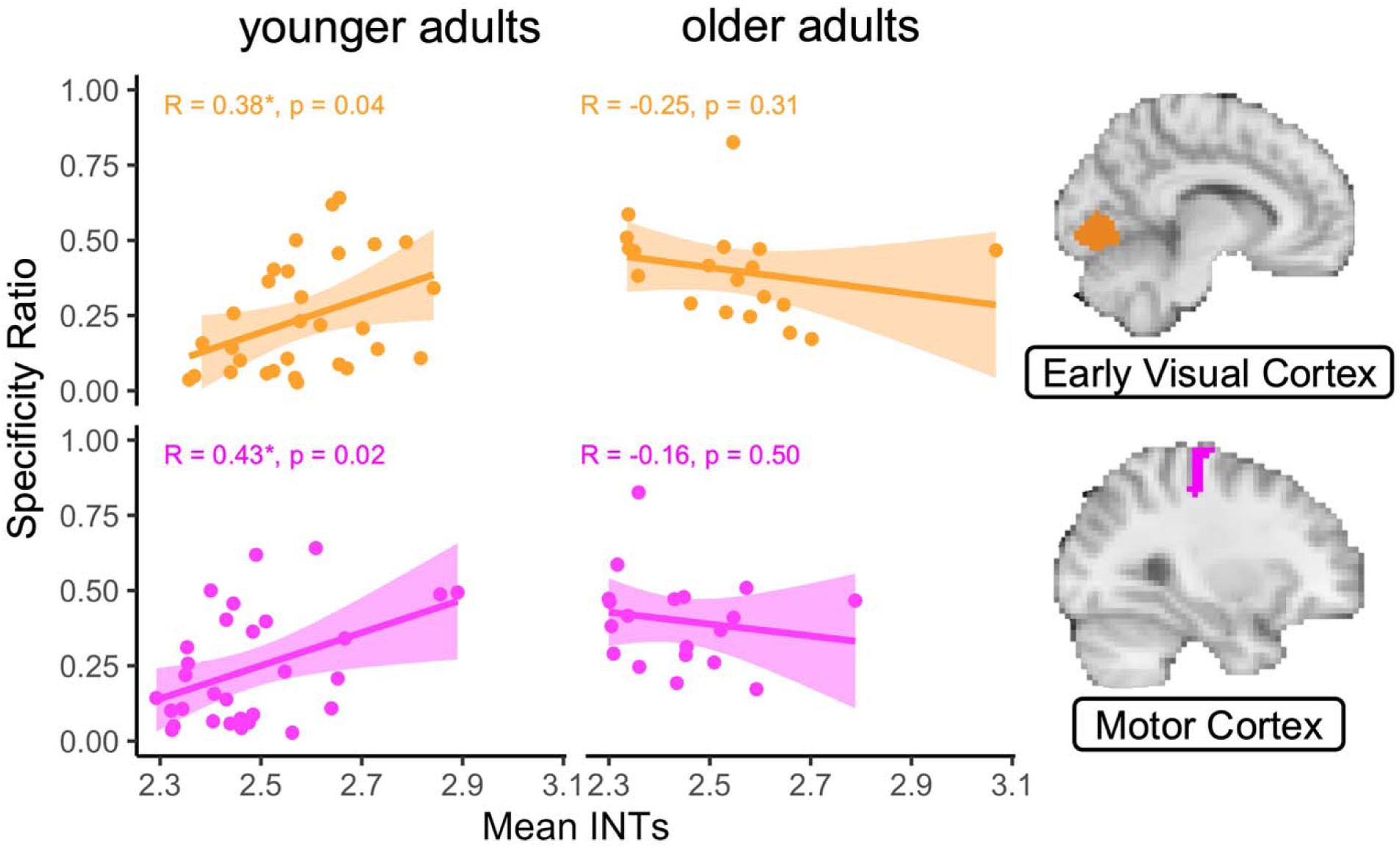
Relationship between INTs and recall specificity in cortical ROIs. Specificity ratio was correlated with INTs in the bilateral early visual cortex (orange) and motor cortex (pink). For the specificity ratio, higher values indicate more gist-like recall and lower values suggest more detailed recall. Each point represents an individual participant. Solid lines indicate the linear regression fit; shaded ribbons indicate 95% CIs. In younger adults, there was a positive correlation between specificity ratio and INTs in the early visual and motor cortex, however these correlations did not survive FDR multiple comparisons correction. There were no significant correlations between specificity ratio and INTs in the early visual or motor cortex in older adults.

We found a positive correlation between mean INTs and specificity scores in the early visual cortex and motor cortex of younger adults, but these did not survive correction for multiple comparisons (Early Visual cortex: R = 0.38, p = 0.04, p_corrected_ = 0.11; Motor cortex: R = 0.43, p = 0.02, p_corrected_ = 0.10). In older adults there were no significant correlations in early visual cortex or motor cortex (ps > 0.31, ps_corrected_ > 0.46). In sum, across several cortical ROIs, we observed a relationship between INTs and memory specificity in younger adults, suggesting that the rate of change in the neural signal across the brain may relate to the specificity of encoding and is not restricted to the hippocampus.

### Relationship between neuropsychological tests, hippocampal INTs, and specificity of memory

We did not have *a priori* predictions about whether INTs would be related to measures of cognitive impairment in older adults. We did not have neuropsychological measures for everyone who completed the experiment; however, we had the complete battery of test scores from 19 older adults. We conducted an exploratory analysis to see if there were any correlations between hippocampal INTs in the anterior-medial and posterior-lateral cluster with the neuropsychological test scores from older adults. We also correlated recall specificity scores with the neuropsychological test scores. We did not find any significant relationship between hippocampal INTs and neuropsychological test scores (Supplemental Table S3) and no significant relationship between specificity ratio and neuropsychological test scores (Supplemental Table S4).

## Discussion

The present study examined age-related differences in hippocampal neural timescales during naturalistic movie-viewing and their relationship to memory specificity. We first replicated the canonical anterior-to-posterior hippocampal timescale gradient in younger adults during naturalistic movie-viewing. Second, older adults exhibited a striking *reversal* of this gradient, with longer timescales in the posterior-lateral hippocampus relative to the anterior-medial hippocampus. Finally, longer neural timescales in the posterior-lateral hippocampus predicted more gist-like recall in younger adults. Strikingly, this brain-behavior relationship also extended to cortical regions including early visual and motor cortex, suggesting that neural timescales may be a signature of a brain-wide encoding state rather than a hippocampus-exclusive mechanism. Together, these findings demonstrate that aging is associated with reorganization of hippocampal temporal dynamics and suggest that hippocampal and cortical neural timescales during encoding may be related to later memory specificity.

This study extends previous findings of hippocampal neural timescale gradients during rest and spatial navigation tasks (Bouffard et al., 2023; Brunec et al., 2018; Raut et al., 2020) to naturalistic movie-viewing. Our findings establish that the timescale gradient is a robust, stable property of hippocampal organization that is observed across tasks in younger adults. In older adults, however, we observed a *reversal* of the gradient. This could reflect a profound reorganization of the temporal dynamics underlying hippocampal function. Critically, this reversal was observed alongside by a shift toward more gist-like memory in older adults, consistent with the decrease in memory specificity seen in aging (Delarazan et al., 2023; Greene & Naveh-Benjamin, 2020, 2024; Grilli & Sheldon, 2022; Levine et al., 2002; Tinner, 2026).

One possibility is that the reversed gradient is driven by a slowing of temporal dynamics in the posterior hippocampus, relative to the anterior hippocampus. If fast-changing signals in the posterior hippocampus are necessary for capturing rapidly unfolding, fine-grained information (Evensmoen et al., 2015; Honey et al., 2012; Paz et al., 2010; Rait et al., 2025), then a reduction in signal updating speed in this region may impair the encoding of event-specific details, shifting memory toward coarser, gist-level representations. A caveat, though, is that we did not find a direct relationship between timescales in the hippocampus and memory specificity in older adults.

Another possibility is that the change in the gradient could reflect brain-wide changes in aging, including interactions between brain areas. There is evidence that aging differentially affects connectivity along the hippocampal long axis. For example, Damoiseaux et al. (2016) reported that advanced age was associated with lower functional connectivity between posterior (but not anterior) hippocampus and default mode network regions, as well as reduced interhemispheric hippocampal connectivity. There may be accompanying changes in the neural timescales in the posterior relative to the anterior hippocampus. Alternatively, Stark et al. (2021) found that aging was associated with a pronounced reduction in functional connectivity between the anterior hippocampus and parahippocampal cortex, while posterior connectivity was less affected. This could reduce the capacity of the anterior hippocampus to maintain stable, longer timescales relative to the posterior hippocampus, which characterizes its function in younger adults. Another suggestive piece of evidence comes from recent inhibitory transcranial magnetic stimulation evidence, which suggests a causal relationship between hippocampal functional connectivity and neural timescales (Coughlan et al. 2023).

The observed age-related gradient reversal might also fit within a broader developmental arc of hippocampal timescale organization. Varga et al. (2025) examined resting-state fMRI and task-based fMRI in a large sample spanning ages 5 to 34 years and found that young children exhibit a pattern that closely mirrors what we observed in older adults: posterior hippocampal timescales greater than anterior timescales. Across development, anterior hippocampal autocorrelation increased linearly, ultimately surpassing posterior autocorrelation before age 30, demonstrating a gradient reversal in which children start with longer timescales in the posterior but by adulthood the anterior hippocampus has longer timescales than the posterior. Similar phenomena were observed in fMRI data during a spatial navigation task comparing children and young adults. Crucially, Varga et al. found that longer neural timescales in the anterior hippocampus were related to increased integration during navigation, converging with past work (Bouffard et al., 2023).

An unexpected finding was the relationship between neural timescales and memory specificity in younger adults. While our hypothesis were based on theories that timescales anterior hippocampus are related to gist representations and that timescales in posterior hippocampus are related to detailed representations (Evensmoen et al., 2015; Poppenk et al., 2013b; Strange et al., 2014), we did not, have strong predictions about brain-behavior relationships. One possibility was that when posterior hippocampal signals slow down, the encoding of fine-grained details is compromised, leading to more gist-like recall and a positive relationship between timescale and specificity ratio. Alternatively, if anterior timescales shorten, this could disrupt the maintenance of the stable event representations during encoding, leading to poorer recall of the episode, resulting in more gist-like recall and a negative relationship between timescales and specificity ratio. This is consistent with recent findings demonstrating that stable, slowly drifting hippocampal signals facilitate the formation of temporally organized memory representations (Rait et al., 2025). These two possibilities make opposite predictions for the direction of the relationship between the timescales and specificity ratio in the anterior vs posterior hippocampus. In fact, we found a trending (albeit not statistically significant) positive relationship between timescales and specificity ratio throughout the whole hippocampus, where longer timescales were related to more gist-like recall in both anterior and posterior hippocampus. This is consistent with the account that a slowing down of signal, across the whole hippocampus, is what drives the shift towards gist-like memory. We interpret this trend with caution given the weak statistical evidence and small sample size, but we believe it merits investigation by future studies.

In an exploratory analysis, we found that the timescale-memory relationship in younger adults was not specific to the hippocampus and was observed in early visual and motor cortex as well. In all these regions, timescales were longer in individuals whose memories were more gist-like. This suggests that the relationship between timescale and memory encoding may reflect a global brain state during encoding that facilitates either detailed or coarse-grained retention across the cortical hierarchy.

Interestingly we only found the timescale-memory relationship in younger adults, not in older adults. While power limitations cannot be ruled out, it is also possible that the decoupling of timescale and memory specificity in older adults reflects a more fundamental disruption in the organization of temporal dynamics across the cortical hierarchy. Widespread changes in BOLD signal variability have been related to age-related deficits in cognitive performance (Garrett et al., 2011, 2013) and age-related decrease in differentiation of neural state transitions (Lugtmeijer et al., 2025) support this idea that there are age-related changes in temporal dynamics that could be reflected in neural timescales of individual voxels. If the increases in neural variability and decreased neural distinctiveness associated with aging disrupts the coordinated temporal architecture that links hippocampal and neocortical dynamics (Honey et al., 2012), this could explain why the brain-wide encoding state that predicts memory specificity in younger adults is absent or reorganized in older adults. Future work with larger samples will be necessary to distinguish these possibilities.

## Limitations and Future Directions

There are several limitations of the present study. First, the number of usable datasets in the older adult sample was small (N = 19), limiting statistical power to detect brain-behavior relationships in this group. Second, although the movie-viewing paradigm offers allows measurement of neural timescales during continuous, naturalistic encoding, it limits experimental influence over neural dynamics. Future studies can examine shifts in hippocampal dynamics under more controlled conditions.

Future research should also examine the mechanisms underlying the gradient reversal in aging more directly. Combining measures of functional connectivity, neural timescales, and memory specificity within the same participants would help to establish whether age-related changes in connectivity mediate the gradient reversal and its downstream effects on memory. Furthermore, investigations of hippocampal timescales in clinical populations, such as those with mild cognitive impairment or Alzheimer’s disease, can yield insights into how hippocampal dynamics reorganize in the presence of disease, like was shown in cases of temporal lobe epilepsy (Bouffard et al., 2026). Additionally, given the striking parallel between the hippocampal gradient reversal in older adults and a similar gradient change observed in children by Varga et al. (2025), future lifespan studies spanning childhood through older adulthood would be particularly valuable for characterizing the full developmental trajectory of hippocampal timescale organization.

## Conclusions

The present findings demonstrate that aging is associated with a reorganization of hippocampal neural timescale gradients, involving a reversal of the canonical anterior-posterior gradient observed in younger adults. This reversal parallels the well-documented shift toward gist-based memory in aging and may reflect broader age-related changes in hippocampal functional connectivity. Furthermore, the relationship between neural timescales and memory specificity appears to reflect a brain-wide encoding state in younger adults, extending beyond the hippocampus to sensory and motor cortices. These findings advance our understanding and situate hippocampal temporal dynamics within both a lifespan developmental context and a broader hierarchical framework of episodic memory processing in the brain.

## Supplemental materials

**Table S1.** Neuropsychological test scores from older adults (N = 19)

| **Neuropsychological Test** | **Mean** | **SD** | **Range** | **Total Possible** |
| --- | --- | --- | --- | --- |
| *Montreal Cognitive Assessment (MoCA)* | 27.58 | 1.92 | [24,30] | 30 |
| *Craft 21 Recall Immediate (Verbatim)* | 24.89 | 6.21 | [13,36] | 44 |
| *Craft 21 Recall Immediate (Paraphrased)* | 18.63 | 3.62 | [10,24] | 25 |
| *Craft 21 Recall Delayed (Verbatim)* | 22.26 | 7.42 | [7,36] | 44 |
| *Craft 21 Recall Delayed (Paraphrased)* | 17.58 | 4.05 | [8,24] | 25 |
| *Multilingual Naming Test (MINT)* | 31.21 | 1.40 | [28,32] | 32 |

**Table S2.**
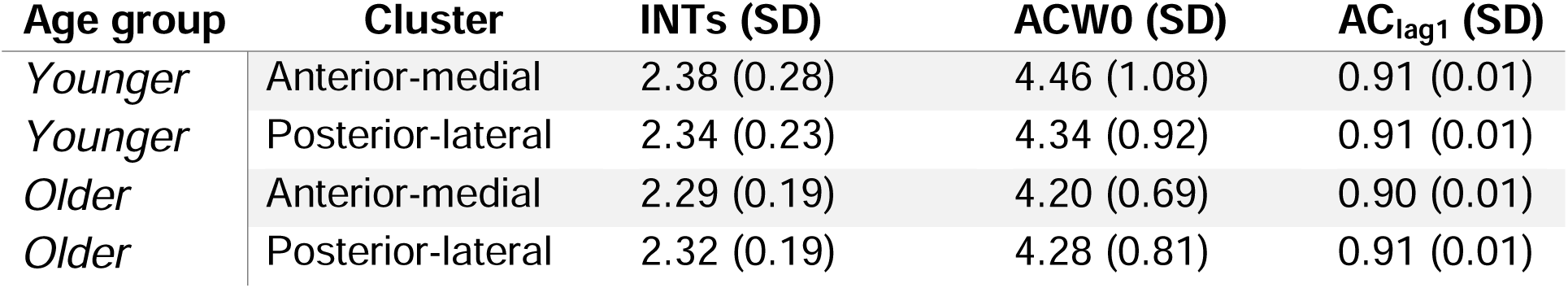
Average intrinsic neural timescale metrics per hippocampal cluster.

| **Age group** | **Cluster** | **INTs (SD)** | **ACW0 (SD)** | **AC<sub>lag1</sub> (SD)** |
| --- | --- | --- | --- | --- |
| *Younger* | Anterior-medial | 2.38 (0.28) | 4.46 (1.08) | 0.91 (0.01) |
| *Younger* | Posterior-lateral | 2.34 (0.23) | 4.34 (0.92) | 0.91 (0.01) |
| *Older* | Anterior-medial | 2.29 (0.19) | 4.20 (0.69) | 0.90 (0.01) |
| *Older* | Posterior-lateral | 2.32 (0.19) | 4.28 (0.81) | 0.91 (0.01) |

**Table S3.** Pearson’s correlation between neuropsychological test scores and INTs in the Anterior-medial and Posterior-lateral hippocampal clusters.

| Neuropsych measure | Anterior-medial Cluster |  |  |  | Posterior-lateral Cluster |  |  |  |
| --- | --- | --- | --- | --- | --- | --- | --- | --- |
|  | r | 0.95% CI | p | p_fdr | r | 0.95% CI | p | p_fdr |
| *Montreal Cognitive Assessment (MoCA)* | 0.01 | [-0.49, 0.51] | >0.999 | 1 | -0.14 | [-0.60, 0.38] | >0.999 | 1 |
| *Craft 21 Recall Immediate (Verbatim)* | -0.19 | [-0.63, 0.33] | >0.999 | 1 | -0.42 | [-0.76, 0.10] | 0.421 | 1 |
| *Craft 21 Recall Immediate (Paraphrased)* | -0.30 | [-0.70, 0.22] | >0.999 | 1 | -0.46 | [-0.78, 0.04] | 0.356 | 1 |
| *Craft 21 Recall Delayed (Verbatim)* | -0.19 | [-0.63, 0.33] | >0.999 | 1 | -0.37 | [-0.73, 0.15] | 0.465 | 1 |
| *Craft 21 Recall Delayed (Paraphrased)* | -0.35 | [-0.72, 0.18] | >0.999 | 1 | -0.49 | [-0.79, 0.01] | 0.338 | 1 |
| *Multilingual Naming Test (MINT)* | 0.05 | [-0.45, 0.54] | >0.999 | 1 | -0.14 | [-0.59, 0.39] | >0.999 | 1 |

**Table S4.**
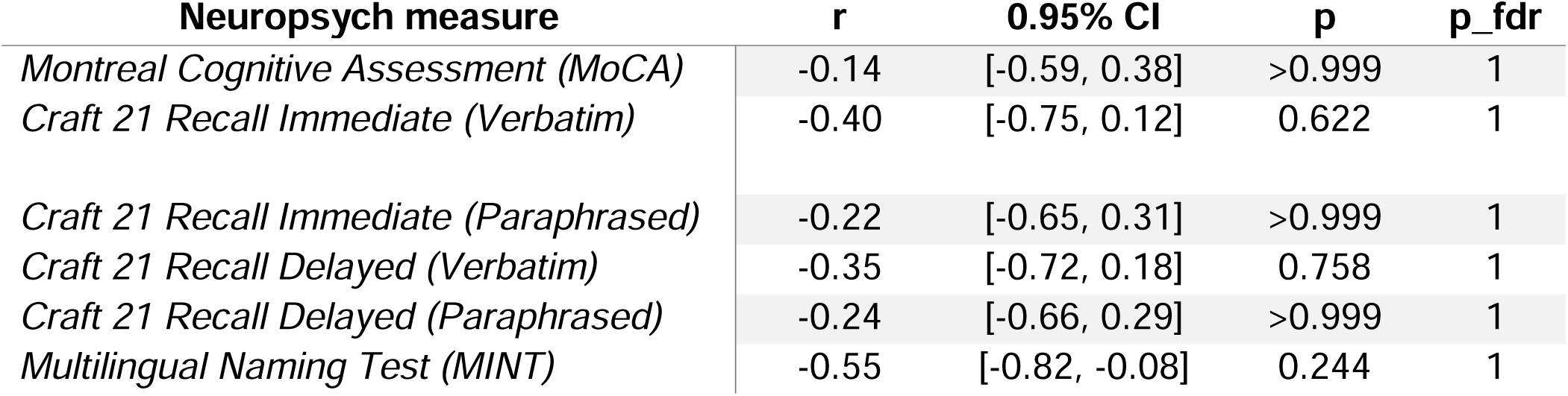
Pearson’s correlation between neuropsychological test scores and specificity ratio.

| **Neuropsych measure** | **r** | **0.95% CI** | **p** | **p_fdr** |
| --- | --- | --- | --- | --- |
| *Montreal Cognitive Assessment (MoCA)* | -0.14 | [-0.59, 0.38] | >0.999 | 1 |
| *Craft 21 Recall Immediate (Verbatim)* | -0.40 | [-0.75, 0.12] | 0.622 | 1 |
| *Craft 21 Recall Immediate (Paraphrased)* | -0.22 | [-0.65, 0.31] | >0.999 | 1 |
| *Craft 21 Recall Delayed (Verbatim)* | -0.35 | [-0.72, 0.18] | 0.758 | 1 |
| *Craft 21 Recall Delayed (Paraphrased)* | -0.24 | [-0.66, 0.29] | >0.999 | 1 |
| *Multilingual Naming Test (MINT)* | -0.55 | [-0.82, -0.08] | 0.244 | 1 |

**Figure S1.**
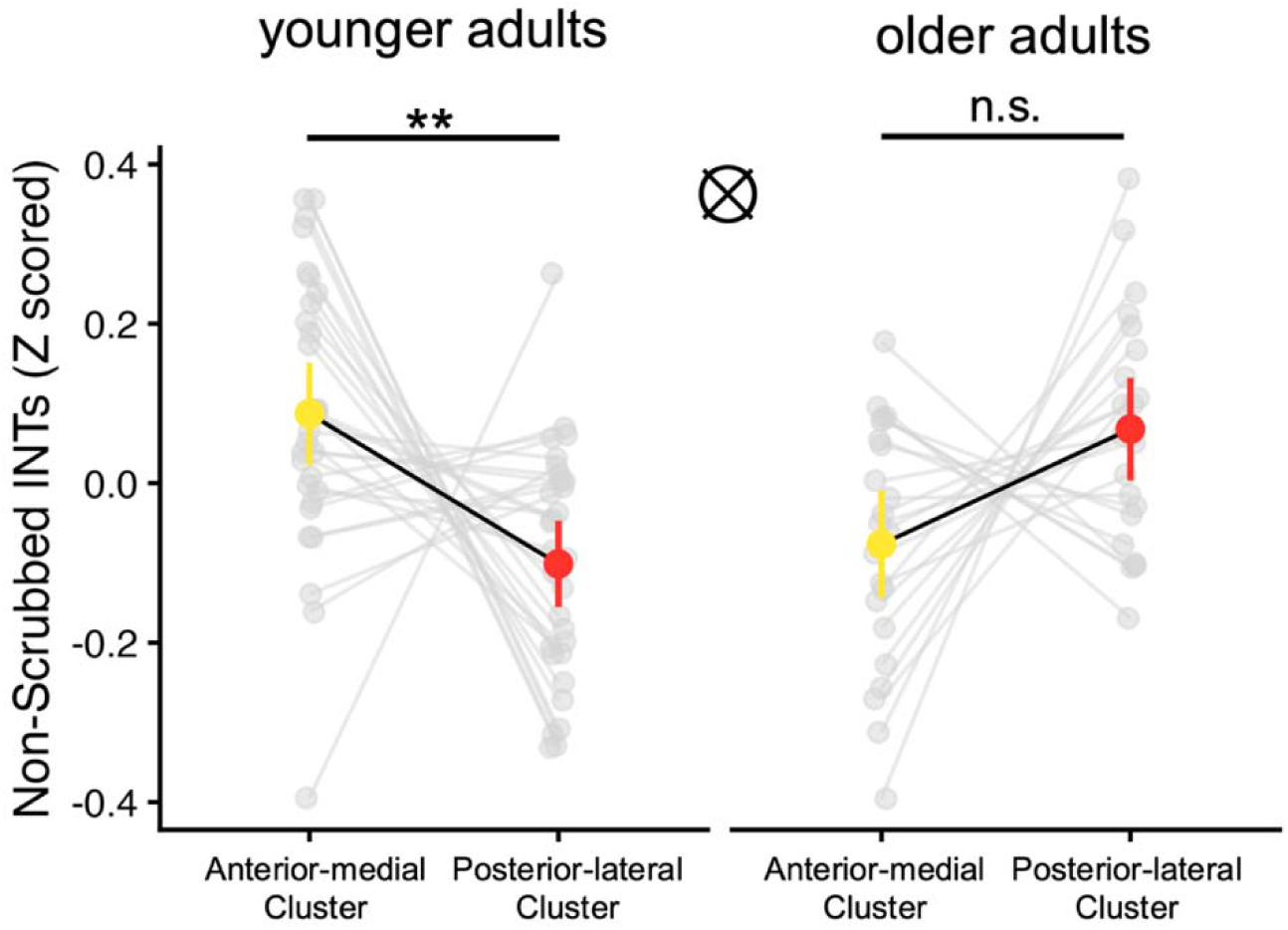
Non-scrubbed hippocampal INTs gradient in younger and older adults. To understand how spike scrubbing affects the INTs measure, we analyzed the non-scrubbed version of the data. All other pre-processing steps were the same, and the same participant inclusion criteria was applied. The only exception was that the motion spike scrubbing step was not applied. We first examine whether the gradient reversal observed in older adults is also found when no scrubbing is applied. We then correlated non-scrubbed INTs with the specificity ratio. We examined whether the INTs computed from the non-scrubbed data was organized in a gradient from the anterior-medial to the posterior-lateral cluster in younger and older adults. We ran a linear mixed effects model on the non-scrubbed INTs for each voxel and included fixed effect predictors for hemisphere (Left, Right), age group (Younger, Older), and cluster (anterior-medial, posterior-lateral). The full model formula: INTs_non-scrubbed_ ∼ hemisphere * age group * cluster + (cluster|participant). For post hoc tests of the main effects and interactions, the degrees of freedom was calculated using an asymptotic method (reporting z ratio instead of t statistic) and p values were adjusted for multiple comparisons using the Tukey method. We did not find a significant main effect of hemisphere, age group, or cluster. We found a significant interaction between age group and cluster (F(1,50.01) = 13.73, p < 0.001). Analysis of the interaction between age group and cluster revealed that the INTs in the anterior-medial cluster were greater than the posterior-lateral cluster in younger adults (z = 3.26, p = 0.006). In older adults we found that INTs in the anterior-medial cluster were not significantly different than the posterior-lateral (z = 2.09, p = 0.153). There was a significant interaction between hemisphere and age group (F(1, 12528.00) = 6.97, p = 0.008). Post hoc tests of this interaction suggest there is a nominal difference in INTs between the left and right hemisphere in older adults (z = 2.34, p = 0.09), however this did not reach significance. The average non-scrubbed INTs in the anterior-medial cluster is depicted in yellow and the average non-scrubbed INTs in the posterior-lateral cluster is depicted in red. Error bars represent the 95% CIs, ** p > 0.01.

**Figure S2.**
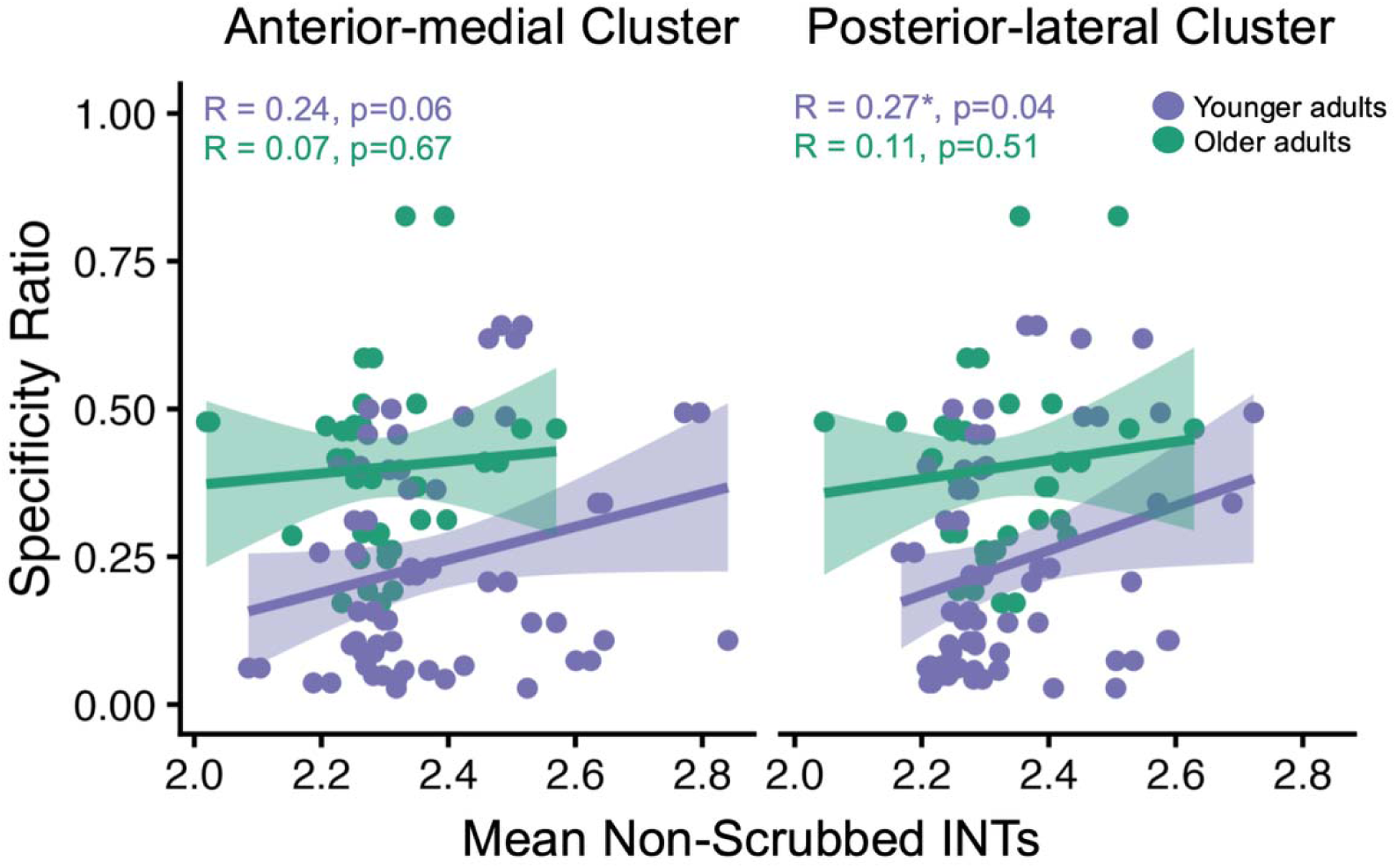
Correlation between specificity ratio and non-scrubbed INTs in the anterior-medial and posterior-lateral clusters. To understand how spike scrubbing affects the relationship between INTs and memory specificity, we correlated the non-scrubbed hippocampal INTs and specificity scores. The INTs was computed from the non-scrubbed data for each voxel in the left and right hippocampus. The non-scrubbed INTs was then averaged within each cluster. Average INTs per cluster was correlated with the specificity scores using Pearson’s correlation. We applied FDR correction for multiple comparisons and report both the uncorrected and uncorrected p values. We found a significant positive correlation between average non-scrubbed INTs and specificity scores in the posterior-lateral cluster in younger adults (Posterior-lateral: R = 0.27, p = 0.04, p_corrected_ = 0.12), however this did not survive multiple comparisons correction. We also found a positive correlation in the anterior-medial cluster, however this did not reach significance (R = 0.24, p = 0.06, p_corrected_ = 0.12). We did not any significant relationships between non-scrubbed INTs and specificity scores in either cluster in older adults (Anterior-medial: R = 0.07, p = 0.67, p_corrected_ = 0.67; Posterior-lateral: R = 0.11, p = 0.51, p_corrected_ = 0.67). When motion spikes are not scrubbed, the positive relationship between average INTs and specificity scores in younger adults still remains, suggesting again that greater hippocampal INTs, or more sluggish signal, during encoding is related to more gist-like recall. When motion spikes are not scrubbed, the positive relationship between average INTs and specificity scores in younger adults remains, although this relationship did not survive multiple comparisons correction. Each point represents the mean non-scrubbed INTs an individual participant and hemisphere. Solid lines indicate the linear regression fit; shaded ribbons indicate 95% CIs.

